# A requirement for Potassium and Calcium Channels during the Endosomal Trafficking of Polyomavirus Virions

**DOI:** 10.1101/814681

**Authors:** Samuel J. Dobson, Jamel Mankouri, Adrian Whitehouse

## Abstract

Following internalisation viruses have to escape the endocytic pathway and deliver their genomes to initiate replication. Members of the *Polyomaviridae* transit through the endolysosomal network and through the endoplasmic reticulum (ER), from which heavily degraded capsids escape into the cytoplasm prior to nuclear entry. Acidification of endosomes and ER entry are essential in the lifecycle of polyomaviruses, however many mechanistic requirements are yet to be elucidated. Alteration of endocytic pH relies upon the activity of ion channels. Using two polyomaviruses with differing capsid architecture, namely Simian virus 40 (SV40) and Merkel cell polyomavirus (MCPyV), we firstly describe methods to rapidly quantify infection using an IncuCyte ZOOM instrument, prior to investigating the role of K^+^ and Ca^2+^ channels during early stages of infection. Broad spectrum inhibitors identified that MCPyV, but not SV40, is sensitive to K^+^ channel modulation. In contrast, both viruses are restricted by the broad spectrum Ca^2+^ channel inhibitor verapamil, however specific targeting of transient or long lasting Ca^2+^ channel subfamilies had no detrimental effect. Further investigation revealed that tetrandrine blockage of two-pore channels (TPCs), the activity of which is essential for endolysosomal-ER fusion, ablates infectivity of both MCPyV and SV40 by preventing disassembly of the capsid, which is required for the exposure of minor capsid protein nuclear signals necessary for nuclear transport. This study therefore identifies a novel target to restrict the entry of polyomaviruses.

**IMPORTANCE:** Polyomaviruses establish ubiquitous, asymptomatic infection in their host. However, in the immunocompromised these viruses can cause a range of potentially fatal diseases. Through the use of SV40 and MCPyV, two polyomaviruses with different capsid organisation, we have investigated the role of ion channels during infection. Here, we show that Ca^2+^ channel activity is essential for both polyomaviruses and that MCPyV is also sensitive to K^+^ channel blockage, highlighting new mechanistic requirements of ion channels during polyomavirus infection. In particular, tetrandrine blockage of endolysosomal-ER fusion is highlighted as an essential modulator of both SV40 and MCPyV. Given that the role of ion channels in disease have been well characterised, there is a large panel of clinically available therapeutics that could be repurposed to restrict persistent polyomavirus infection and may ultimately prevent polyomavirus-associated disease.

## INTRODUCTION

Polyomaviruses (PyVs) are small double stranded DNA viruses that establish persistent infections in their hosts. Whilst infections are generally asymptomatic, PyVs can cause severe disease in the immunosuppressed. Common examples include BKPyV-associated nephropathy and haemorrhagic cystitis, and JCPyV-induced progressive multifocal leukoencephalopathy (PML) (1–3).

In ∼80% of Merkel cell carcinoma (MCC) cases, Merkel cell PyV (MCPyV) infection, clonal integration and UV-mediated mutation of the viral genome occur prior to tumour cell expansion, with truncation of the large tumour antigen (LT) rendering MCPyV replication defective (4–7). Both MCPyV small T and truncated LT proteins are required for MCC survival and proliferation (8–10). Although the continuous infection of MCPyV is not implicated in MCC, the routes of virus entry into susceptible cells remain elusive. The capsids of all PyVs consist of 72 VP1 pentamers that form an icosahedral structure with T=7d symmetry and mediate initial surface receptor binding (11–13). Under each pentamer sits a minor capsid protein linking VP1 to the viral genome (11). The majority of PyVs, including SV40, BKPyV and JCPyV encode two minor capsid proteins (VP2 and VP3) which are incorporated into the capsid. MCPyV is however part of a small clade of PyVs that only express one minor capsid protein (VP2) (14).

All PyVs must deliver their genomes to the nucleus, commonly achieved by trafficking through the endosomal system (15, 16). Initial attachment varies across PyV species but typically involves sialylated glycans. SV40 interacts with MHC-1 and GM1 gangliosides in lipid rafts, whilst MCPyV interacts with sulphated glycosaminoglycans including heparan sulphate or chondroitin sulphate prior to secondary interactions with sialylated glycans to facilitate virus penetration (17–21). Following binding, JCPyV enters cells through clathrin-mediated endocytosis, whilst SV40, MCPyV and BKPyV enter via caveolar/lipid rafts (22–27). virions traffic through the endosomal system and in response to endosomal cues, including endosome acidification, initiate proteolytic rearrangements of the capsid prior to retrograde trafficking to the endoplasmic reticulum (ER) (25, 28–30). Within the ER, virions are further disassembled, exposing nuclear localisation signals (NLSs) that transport capsids to the nucleus via importins (31–37). Despite this knowledge, the endosomal cues that permit PyV trafficking remain poorly understood.

Emerging studies suggest that the current description of virus entry processes involving acidification alone are too simplistic and that the accumulation of other ions including K^+^ and Ca^2+^ influence virus trafficking (38–43). In the context of PyV infection, Ca^2+^ ions have been shown to affect the structure and organisation of virus particles, regulating their disassembly through virion swelling (40, 44–46). However, despite the evidence that cellular ion channels are targeted by a wide range of viruses to enhance specific lifecycle stages, their role during PyV entry has not been defined (40, 41, 53, 42, 43, 47–52).

In this study, we used two distantly related PyVs, namely SV40 and MCPyV, to determine if K^+^ or Ca^2+^ channels are required for viral progression through the endosomal system. To achieve this, reporter-containing MCPyV pseudovirions (PsVs) that behave in a similar manner to WT viruses were used alongside native SV40 virions to specifically assess virus entry through high-throughput fluorescence-based detection systems. Herein, we show that MCPyV and SV40 are differentially sensitive to K^+^ channels and transient (T-type) Ca^2+^ channel inhibitors. We further identify a shared requirement of SV40 and MCPyV for the activity of endosomal nicotinic acid adenine dinucleotide phosphate (NAADP)-Sensitive Two-Pore Ca^2+^ Channels (TPCs) that regulate ER-endosome membrane contact sites. These findings reveal potential therapeutic drug targets for PyVs and enhance our understanding of the virus entry processes.

## RESULTS

### Fluorescence-based detection of SV40 and MCPyV

A challenge in PyV studies are the limited experimental systems to assess the complete virus lifecycle. We initially established a high throughput fluorescence detection system to monitor the expression of SV40 T-antigens following virus infection using an Incucyte ZOOM instrument, in a manner comparable to previous methods described for Hepatitis C virus measurement (**Fig. 1A-B**) (54, 55). Cells infected with SV40 were fixed, then permeabilised and immunostained to detect T-antigens, prior to automated imaging and analysis to determine the number of T-antigen positive cells. The system could reproducibly quantify the number of SV40 T-antigen positive cells and as such could be used as a rapid method to assess the levels of virus infection (**Fig. 1C-D**). The study of MCPyV is more challenging due to a lack of reproducible infectious systems. We therefore applied reporter-containing PsVs that permit the assessment of MCPyV entry and genome release into the nucleus. MCPyV PsVs transduced target cells more slowly than SV40 infection, with detectable fluorescence observed between 48 to 72 hours post-transduction (hpt), consistent with previously reported timescales **(Fig. 1E-F)** (25). Using these systems, SV40 infections and MCPyV transductions could be performed in a 96- or 24-well plate format, respectively, providing a platform for high-throughput antiviral compound screening.

**Figure 1:**
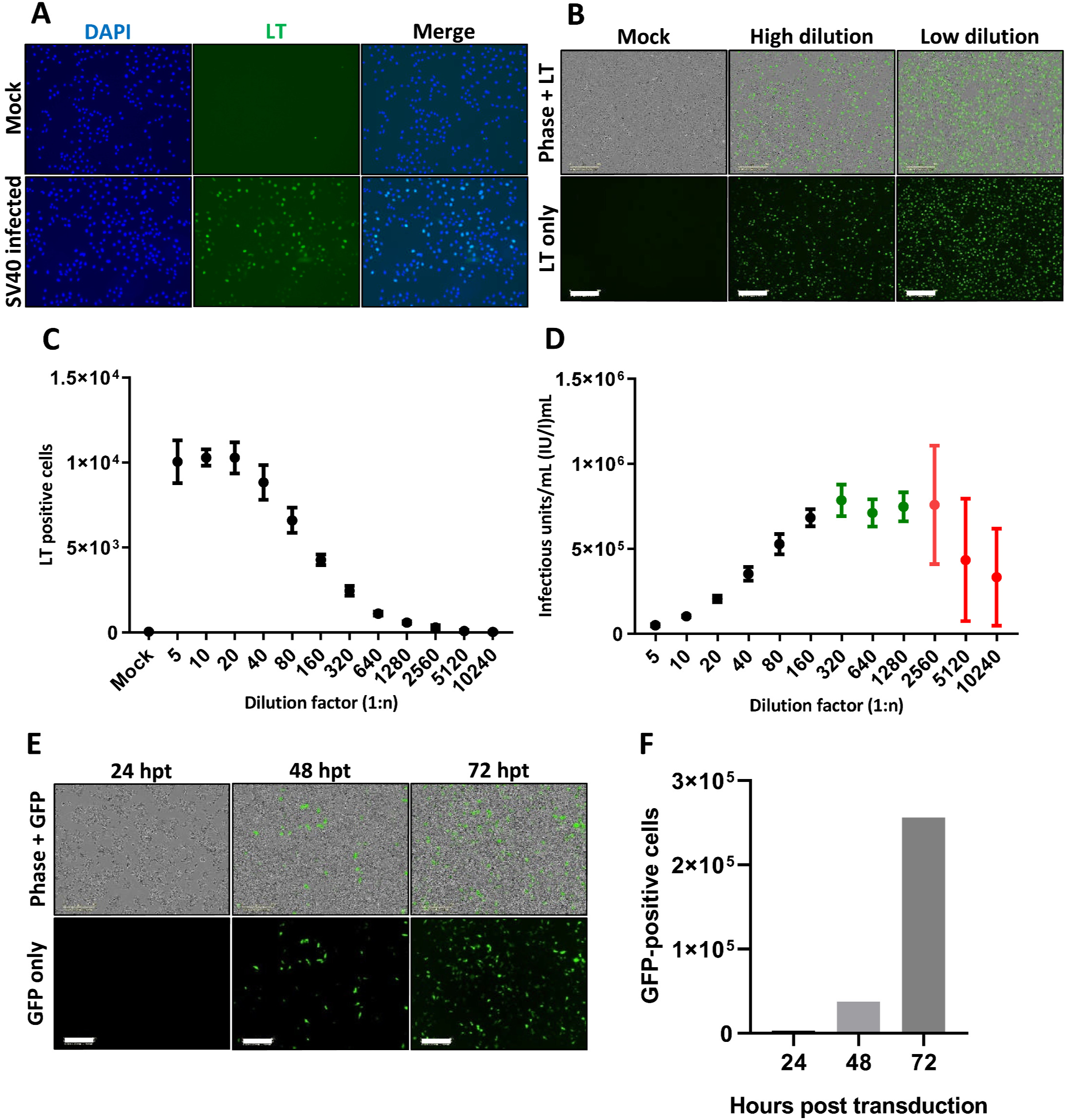
Development of immunofluorescence-based systems to determine SV40 titre and study early events in the lifecycles of SV40 and MCPyV. Visualisation of SV40 infected Vero cells 24 hours post infection by light microscopy (**A**) and Incucyte ZOOM instrument (**B**). For light microscopy DAPI was used to stain nucleic acids whilst SV40 LT/ST specific primary and Alexa Fluor 488 secondary antibodies were used to visualise infected cells. Images were taken using an EVOS II microscope. For validation by Incucyte detection, SV40 infected Vero cells were similarly immunostained before imaging. Shown are representative images using differentially diluted virus stock, scale bar 300 µM. (**C**) SV40 T-antigen positive cell counts were determined by Incucyte ZOOM detection at a range of dilutions. Reversal of dilution factors was performed to determine infectious units/mL (**D**), whereby the plateau represented viral titre (green) and hypervariable replicates were indicative of loss of assay sensitivity (red). (**E**) Viability of Incucyte detection for GFP-expressing MCPyV PsVs was also confirmed by imaging at 24-, 48- and 72-hours post transduction, with autonomous quantification of GFP-positive cells (**F**). Scale bar 300 µM.

It is well established that PyVs traffic through the endo/lysosomal system where acidification initiates proteolytic rearrangements to promote virus disassembly. To validate our Incucyte-based system, we assessed virus infection in the presence of ammonium chloride (NH_4_Cl), a known inhibitor of endosomal acidification. Consistent with previous studies, NH_4_Cl treatment reduced MCPyV and SV40 infection by 87% and 54% respectively, further validating the system (**Fig. 2A & C**). However, despite knowledge that acidification is important, the endosomal progression of PyVs prior to ER translocation remain unclear. To determine whether the viruses enter late endosomes and/or lysosomes, cells were treated with 2-[(4-Bromophenyl) methylene]-N-(2, 6-dimethylphenyl)-hydrazinecarboxamide (EGA) to inhibit lysosomal clustering prior to virus infection. Treatment with EGA led to a 75% and 78% decrease in MCPyV and SV40 infected cells respectively, suggesting that both viruses transverse the late endosome/lysosomal system (**Fig. 2B & D**).

**Figure 2:**
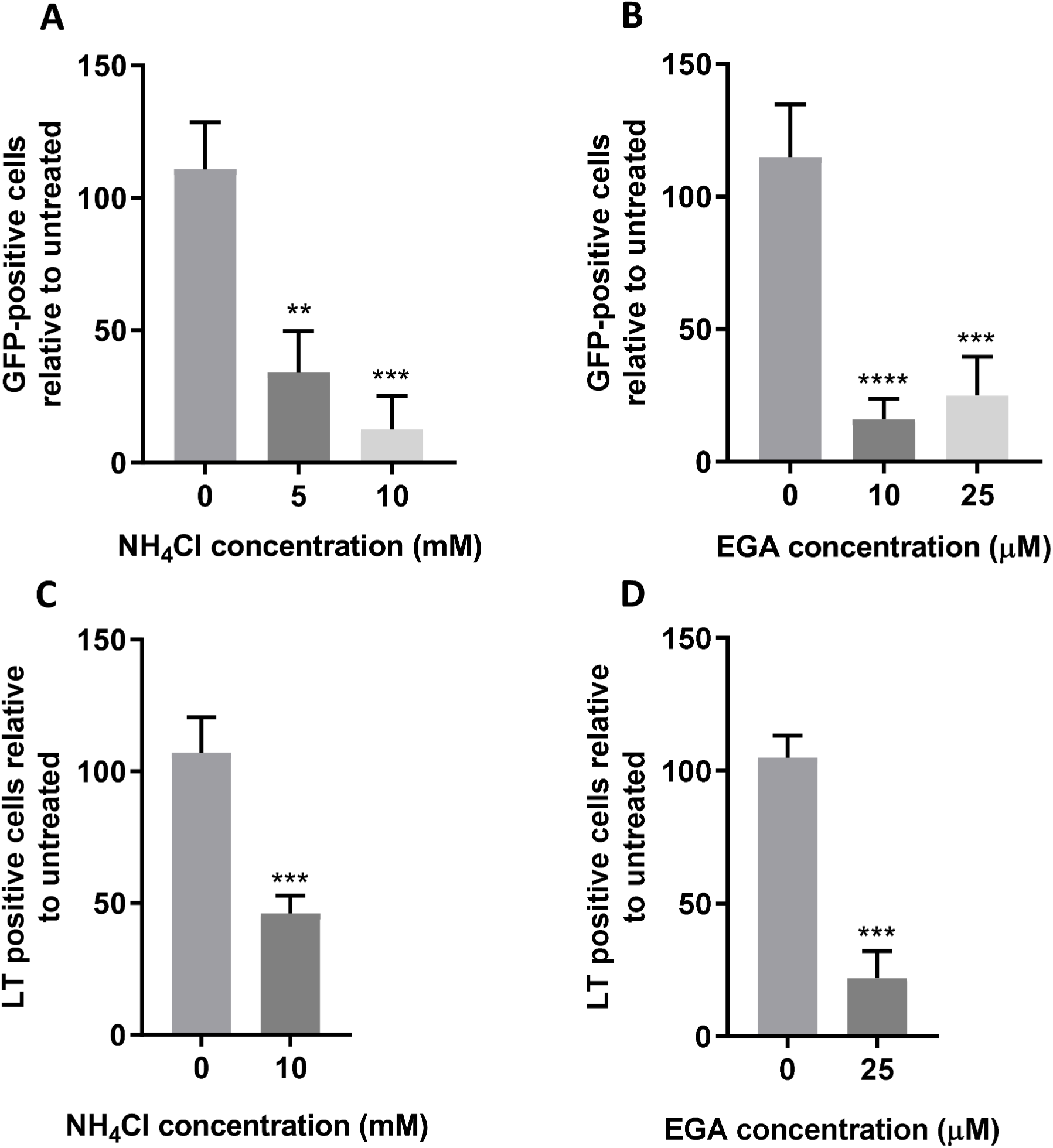
MCPyV and SV40 both enter into acidified endosomes. (**A+B**) 293TT cells were incubated with drug as described for 1 hour before addition of 10 ng VP1-equivalent MCPyV GFP PsVs for 2 hours with occasional agitation. PsV containing medium was removed and replaced with fresh drug-containing medium, with Incucyte detection 72 hours post transduction to determine the number of GFP-positive cells. (**C+D**) Vero cells were incubated with drug as described for 1 hour before addition of SV40 virions at an MOI of 1 for 2 hours with occasional agitation. Fresh drug-containing medium was then added for an incubation of 24 hours before fixation and permeabilisation. SV40 T-antigens were immunostained using an SV40 LT/ST antibody and species-specific Alexa Fluor 488 secondary antibody. Wells were then imaged using an Incucyte ZOOM instrument to determine the number of T-antigen positive cells.

### K^+^ and Ca^2+^ channel inhibition restricts MCPyV entry

Ion channels have emerged as key regulators of virus entry processes. Examples include the negative sense RNA viruses, bunyamwera virus and influenza virus that require K^+^ during endosomal transit to mediate virus priming and endosomal escape (38, 40, 56). We therefore explored whether regulators of two of the major endosomal ion channel families, namely K^+^ or Ca^2+^ channels, were important for PyV entry. Treatment of cells with the broad spectrum K^+^ channel inhibitor tetraethylammonium (TEA) and broad-spectrum Ca^2+^ channel inhibitor verapamil inhibited MCPyV infection by 62% and 57%, respectively, suggesting that both channel families were important during MCPyV entry (**Fig. 3A-B**). In contrast, TEA had little to no-effect on SV40 infection (**Fig. 3C**), whereas Verapamil treatment caused a 57% loss of SV40 infection (**Fig. 3D**). This suggested a requirement for Ca^2+^ channels is conserved across both viruses.

**Figure 3:**
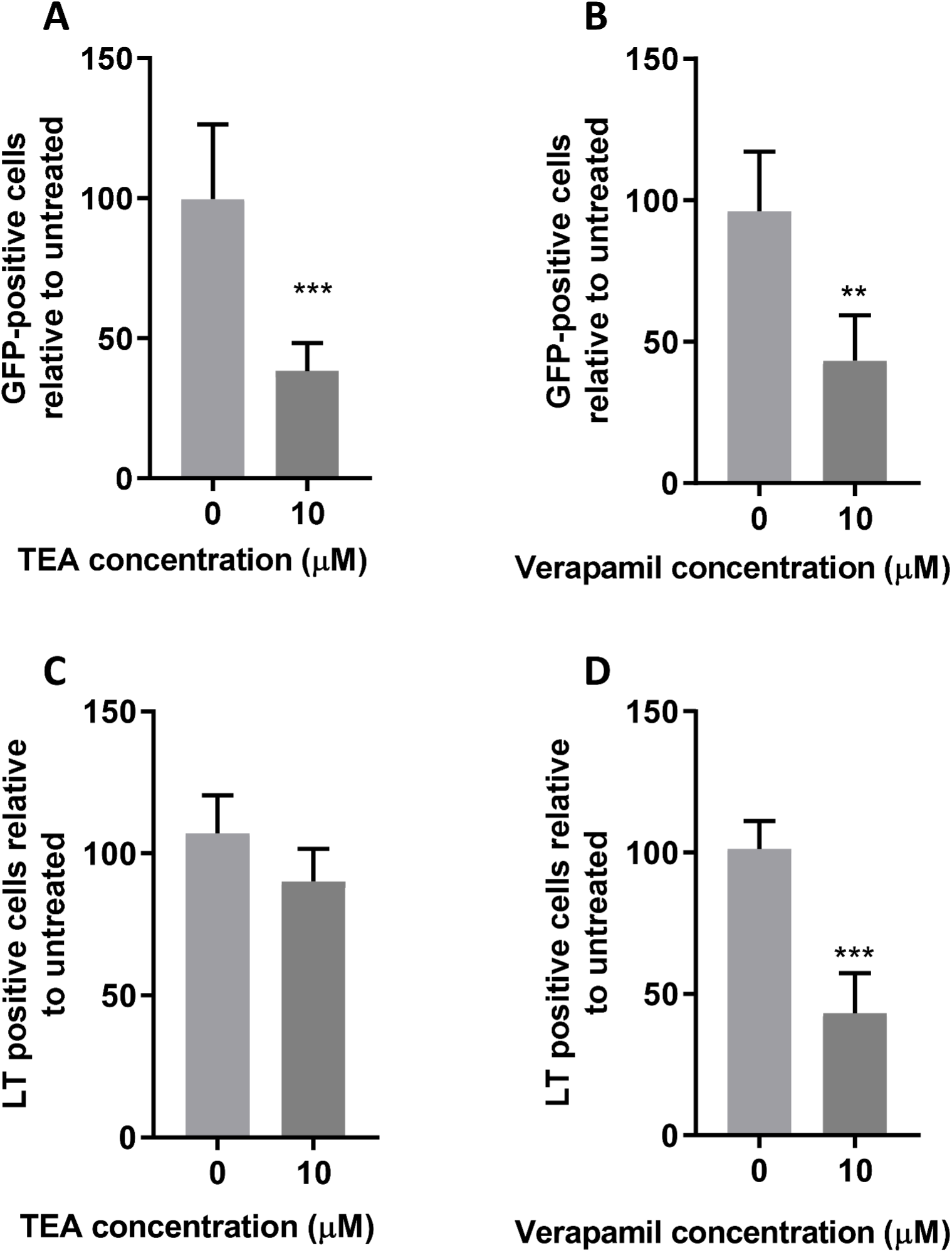
MCPyV and SV40 have a conserved requirement of Ca^2+^ channels, whilst K^+^ channel requirements are only required by MCPyV. (**A+B**) 293TT cells were incubated with drug as described for 1 hour before addition of 10 ng VP1-equivalent MCPyV GFP PsVs for 2 hours with occasional agitation. PsV containing medium was removed and replaced with fresh drug-containing medium, with Incucyte detection 72-hours post transduction to determine the number of GFP-positive cells. (**C+D**) Vero cells were incubated with drug as described for 1 hour before addition of SV40 virions at an MOI of 1 for 2 hours with occasional agitation. Fresh drug-containing medium was then added for an incubation of 24 hours before fixation and permeabilisation. SV40 T-antigens were immunostained using an SV40 LT/ST antibody and species-specific Alexa Fluor 488 secondary antibody. Wells were then imaged using an Incucyte ZOOM instrument to determine the number of T-antigen positive cells.

### Identification of the K^+^ channels required during MCPyV entry

K^+^ channels are the most diverse class of membrane proteins expressed with the cell (57). There are four subfamilies that are ubiquitously expressed across nearly all kingdoms of life: (i) voltage-gated K^+^ channels (K_V_) (6 transmembrane domains (TMDs)), (ii) inwardly rectifying K^+^ channels (K_IR_) (2 TMDs), (iii) tandem pore domain K^+^ channels (K_2P_) (4 TMDs) and (iv) Ca^2+^ activated K^+^ channels (K_Ca_) (6 TMDs) (58). To identify which K^+^ channel subfamilies are required during MCPyV entry, we investigated the effects of KCl (to destroy K^+^ gradients and thus K^+^ channel function) and 4-aminopyridine (4AP, a K_V_ channel blocker) on MCPyV and SV40 infection. Both KCl and 4AP inhibited MCPyV but neither affected SV40 (**Fig. 4A-B**). Furthermore, the anti-malarial drug quinine that promiscuously blocks a variety of K^+^ channels through an unknown mechanism had no effect on either MCPyV or SV40. These data highlighted differences in MCPyV and SV40 entry processes and suggested that MCPyV can be blocked by inhibitors of 4AP sensitive, quinidine insensitive Kv channels.

**Figure 4:**
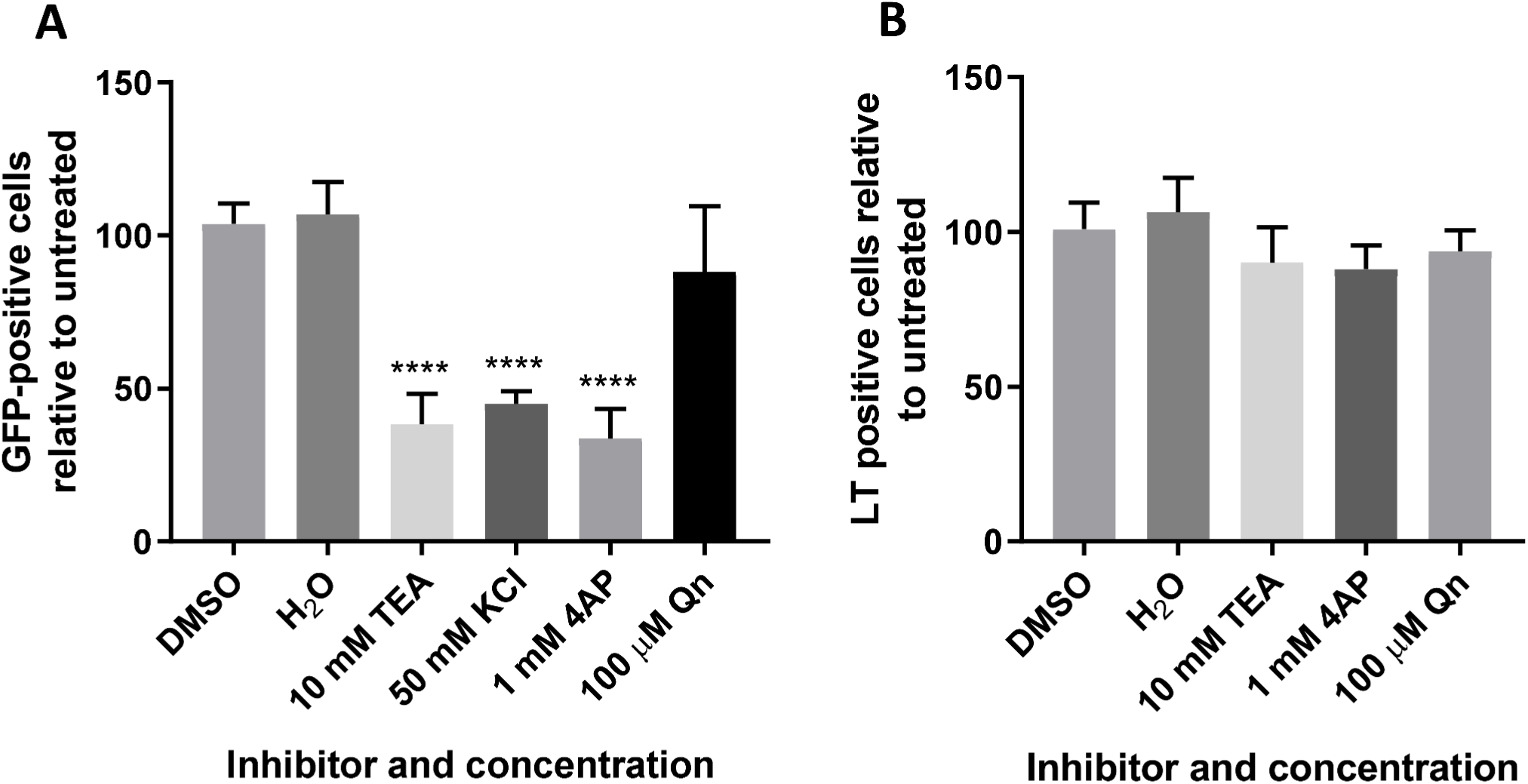
Requirement of K^+^ channel activity during entry is MCPyV specific. (**A**) 293TT cells were incubated with drug as described for 1 hour before addition of 10 ng VP1-equivalent MCPyV GFP PsVs for 2 hours with occasional agitation. PsV containing medium was removed and replaced with fresh drug-containing medium, with Incucyte detection 72 hours post transduction to determine the number of GFP-positive cells. (**B**) Vero cells were incubated with drug as described for 1 hour before addition of SV40 virions at an MOI of 1 for 2 hours with occasional agitation. Fresh drug-containing medium was then added for an incubation of 24 hours before fixation and permeabilisation. SV40 T-antigens were immunostained using an SV40 LT/ST antibody and species-specific Alexa Fluor 488 secondary antibody. Wells were then imaged using an Incucyte ZOOM instrument to determine the number of T-antigen positive cells. 50 mM KCl was omitted due to Vero cell cytotoxicity.

### Blockers of L-type Ca^2+^ channels restrict MCPyV entry

Given that K^+^ channel inhibition did not display a conserved effect upon MCPyV and SV40 entry, the effect of verapamil was further investigated. Verapamil inhibits both Transient (T-type, low-voltage activated) and long lasting (L-type, high-voltage activated) Ca^2+^ channel family members. Therefore, a range of more specific Ca^2+^ blocking drugs were assessed for their effects on SV40 and MCPyV. Treatment with the T-type inhibitor flunarizine led to an 83% inhibition of MCPyV infection, whilst nitrendipine (an L-type Ca^2+^ channel blocker) had no significant effect (**Fig. 5A-B**). Both flunarizine and nitrendipine did not affect SV40 entry suggesting that the requirement for T-type Ca^2+^ channels is limited to MCPyV **(Fig. 5C-D**).

**Figure 5:**
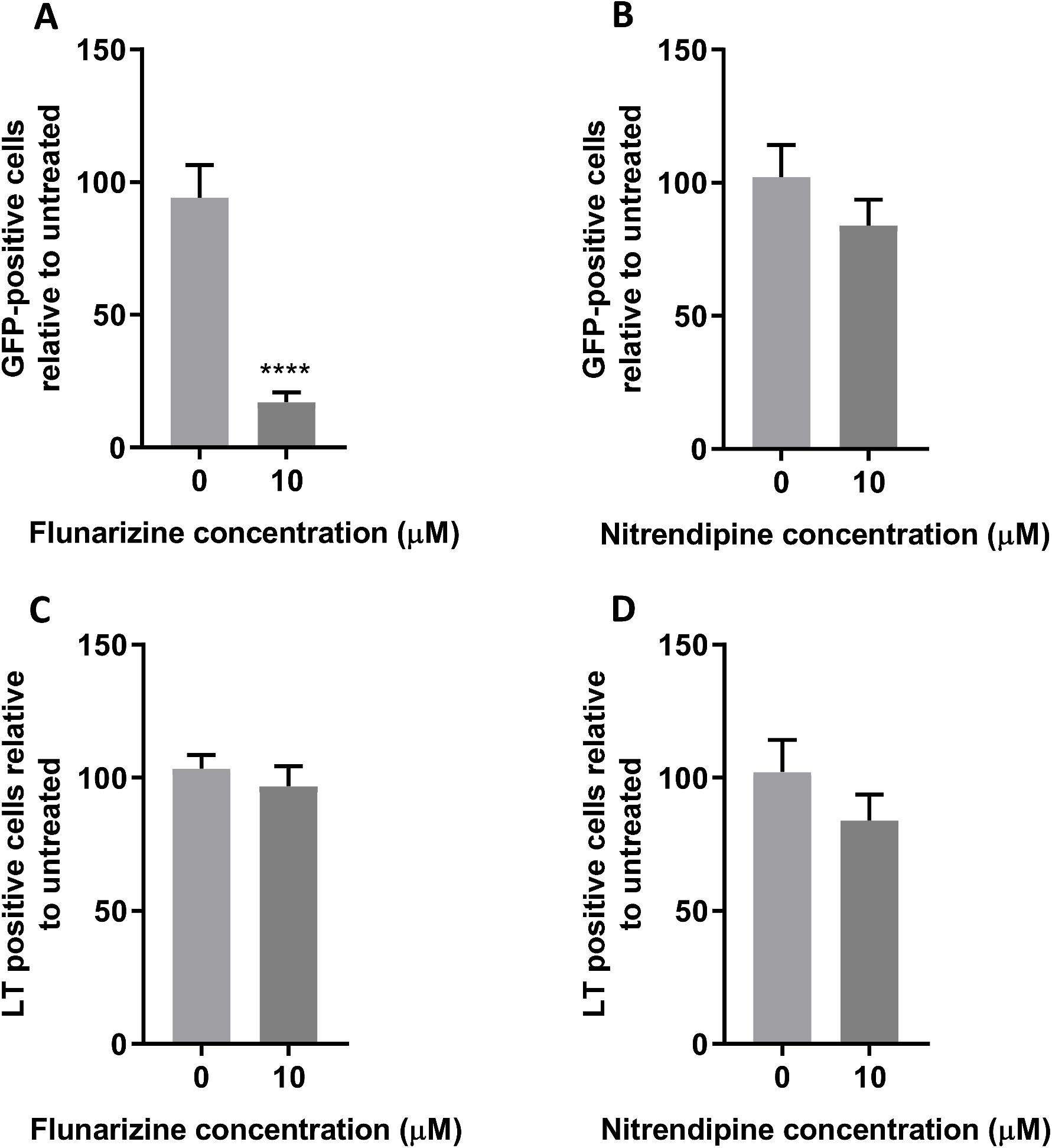
Inhibition of T-type Ca^2+^ channels restricts MCPyV entry, but SV40 is not affected by inhibition of T- or L-type Ca^2+^ channels. (**A+B**) 293TT cells were incubated with drug as described for 1 hour before addition of 10 ng VP1-equivalent MCPyV GFP PsVs for 2 hours with occasional agitation. PsV containing medium was removed and replaced with fresh drug-containing medium, with Incucyte detection 72 hours post transduction to determine the number of GFP-positive cells. (**C+D**) Vero cells were incubated with drug as described for 1 hour before addition of SV40 virions at an MOI of 1 for 2 hours with occasional agitation. Fresh drug-containing medium was then added for an incubation of 24 hours before fixation and permeabilisation. SV40 T-antigens were immunostained using an SV40 LT/ST antibody and species-specific Alexa Fluor 488 secondary antibody. Wells were then imaged using an Incucyte ZOOM instrument to determine the number of T-antigen positive cells.

### Blockers of two pore Ca^2+^ channels inhibit MCPyV and SV40

It has previously been shown that verapamil, alongside a panel of classical L-type inhibitors could inhibit the entry of EBOV (41). Further investigation however identified that EBOV did not not require L-type Ca^2+^ channel activity, with blockage of NAADP-dependent TPCs that regulate endosomal Ca^2+^ signalling, sufficient in preventing endolysosomal fusion of virus-containing vesicles with the ER. Given that PyVs traffic through the ER and verapamil showed an effect that was independent of the assessed Ca^2+^ channel inhibitors, the importance of TPCs during MCPyV and SV40 entry was investigated. Gabapentin, an L-type Ca^2+^ channel inhibitor had no effect on MCPyV or SV40, which was comparable with nitrendipine treatment (**Fig. 6A & C**). However, treatment with the TPC inhibitor tetrandrine led to a striking concentration-dependent inhibition of both viruses, with near complete abolishment of fluorescent cells for MCPyV and SV40 at 5 µM and 10 µM, respectively (**Fig. 6B & C**). Loss of infectivity for both viruses confirmed that NAADP Ca^2+^ channels were essential for PyV infection and may represent a conserved target to restrict a wider range of PyV infections.

**Figure 6:**
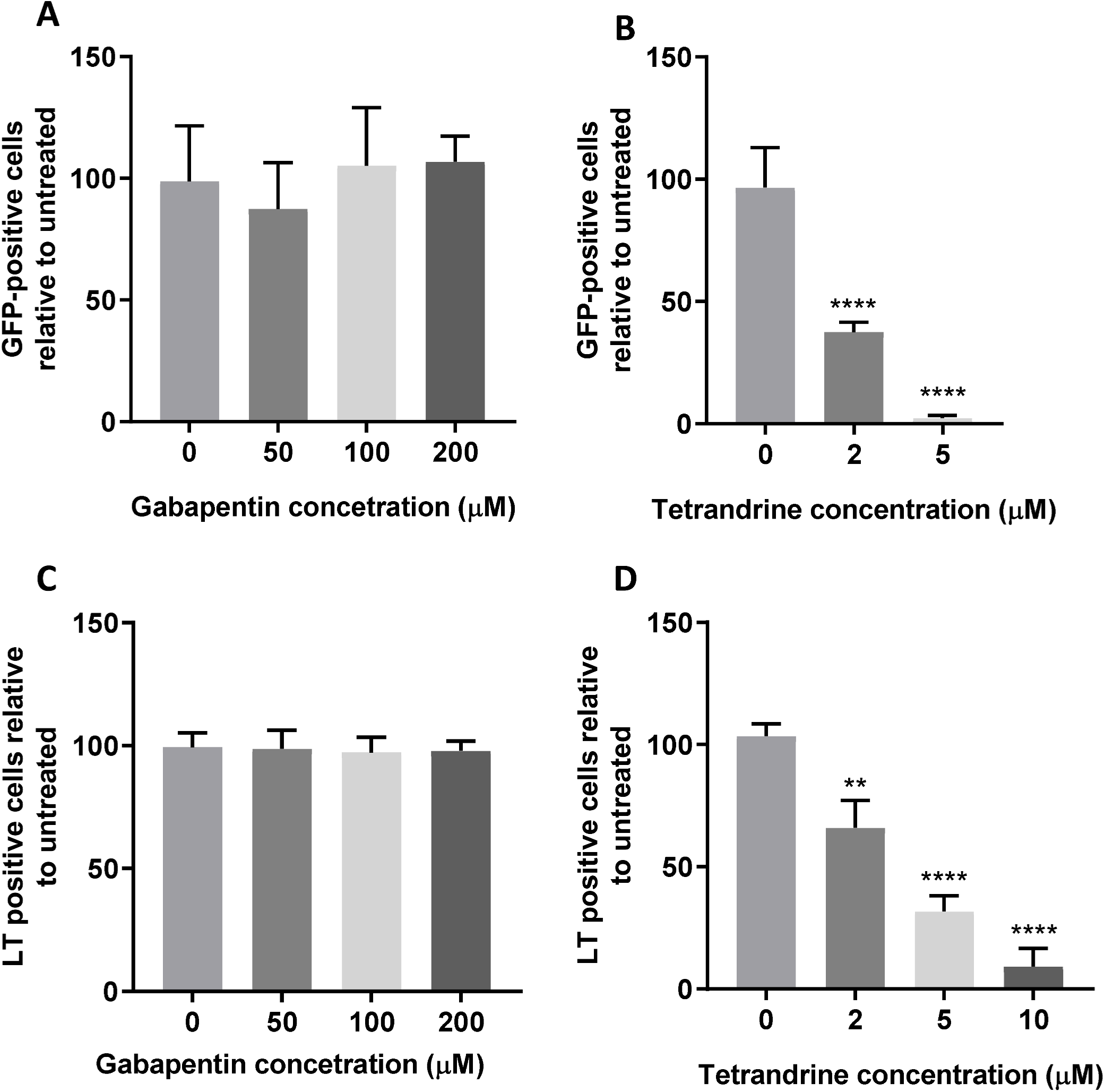
Two pore channel activity is essential for MCPyV and SV40 entry. (**A+B**) 293TT cells were incubated with drug as described for 1 hour before addition of 10 ng VP1-equivalent MCPyV GFP PsVs for 2 hours with occasional agitation. PsV containing medium was removed and replaced with fresh drug-containing medium, with Incucyte detection 72 hours post transduction to determine the number of GFP-positive cells. (**C+D**) Vero cells were incubated with drug as described for 1 hour before addition of SV40 virions at an MOI of 1 for 2 hours with occasional agitation. Fresh drug-containing medium was then added for an incubation of 24 hours before fixation and permeabilisation. SV40 T-antigens were immunostained using an SV40 LT/ST antibody and species-specific Alexa Fluor 488 secondary antibody. Wells were then imaged using an Incucyte ZOOM instrument to determine the number of T-antigen positive cells.

### TPC inhibition prevents SV40 ER disassembly

Although proteolytic rearrangements are initiated in acidifying endosomes, SV40 capsid disassembly sufficient for minor capsid protein exposure does not occur until the virion is processed in the ER (∼6-8 hpi), with detection in the cytoplasm at 10 hpi (30). To confirm a role of TPCs during SV40 entry, virus supernatants were added to cells at 4°C to synchronise infection, prior to the addition of pre-warmed medium containing vehicle or inhibitor for 10 h. Cells were then fixed and immunostained for VP2/3 to detect disassembled virions in the ER and cytoplasm. We observed distinct puncta in cells treated with vehicle or gabapentin (**Fig. 7**). In contrast, cells treated with tetrandrine displayed no detectable puncta confirming that the capsid was unable to disassemble and expose the minor capsid NLSs required for transit to the nucleus. These results highlight an essential requirement for NAADP-stimulated Ca^2+^ channel activity during SV40 infection.

**Figure 7:**
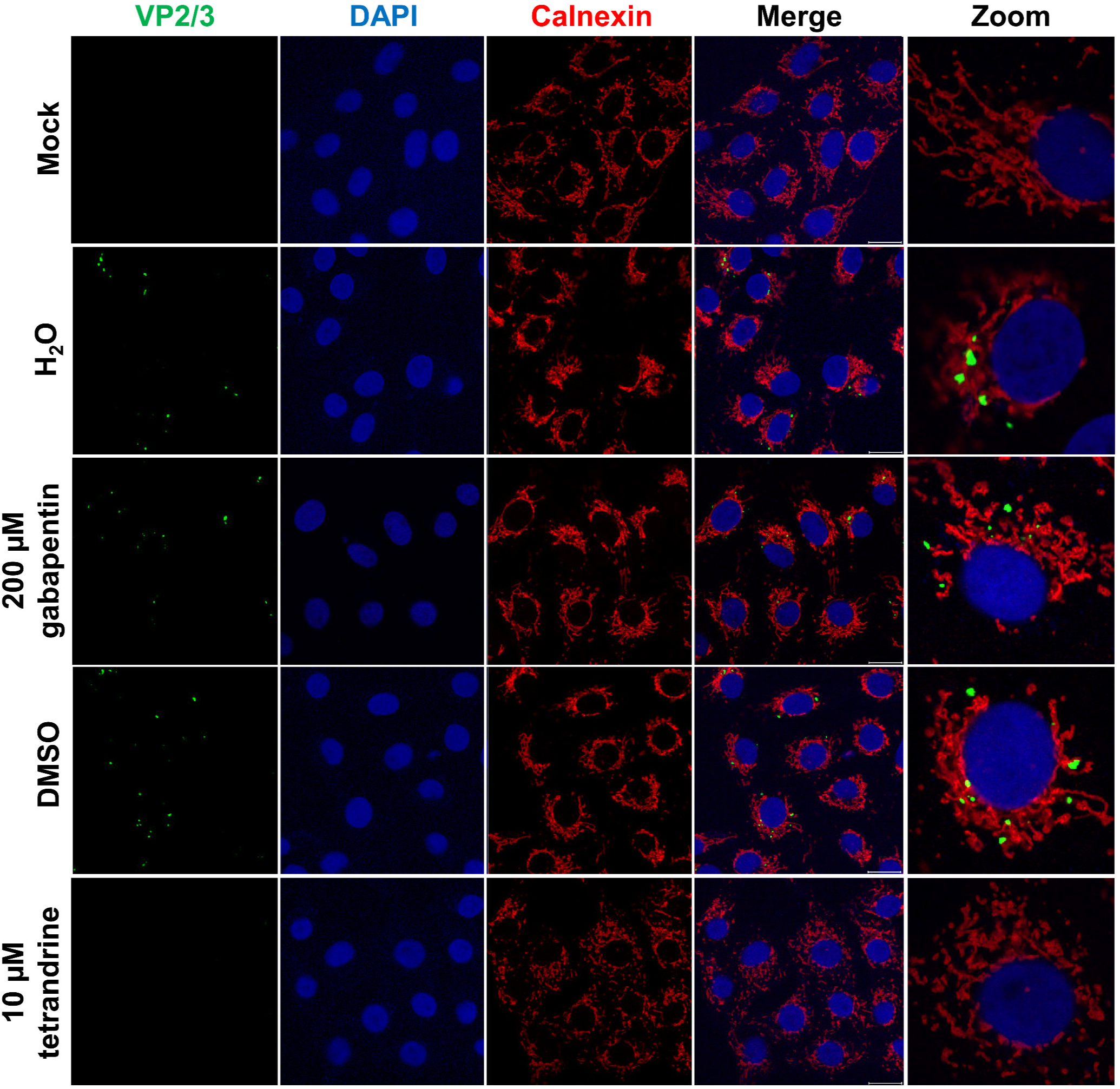
Two pore channel inhibition prevents SV40 disassembly and exposure of minor capsid proteins. Vero cells were chilled at 4 °C for 1 hour before addition of SV40 virions at an MOI of 3 in chilled growth medium. Cells were maintained at 4 °C for 1 hour with occasional agitation to facilitate binding. Pre-warmed growth medium containing drug was then added before incubation at 37 °C for 10 hours before fixation. Following permeabilisation, immunostaining was performed to detect SV40 VP2/3 and the ER using calnexin. DAPI was used to visualise nucleic acids.

## DISCUSSION

To date, studies regarding early events in the lifecycle of PyVs are limited. All studied PyVs traffic through the endo/lysosomal network during virus entry, which we confirmed for both MCPyV and SV40 using newly developed, high-throughput fluorescence-based assays (**Figs. 1-2**) (16). However. the specific routes of endosomal translocation and the host factors required during trafficking to the ER remain largely undefined. Whilst it has long been understood that the acidification of endosomes is essential for PyV entry cues, the endosomal balance of other ions and their crucial roles during the infection of a plethora of viruses is only beginning to emerge (15, 39).

There is a long-standing acceptance that acidification of maturing endosomes and lysosomes is due to the translocation of H^+^, which whilst true, only reflects one aspect of the highly dynamic ionic flux that regulates compartmental pH (59). Given that ion channels regulate a wide variety of cellular functions, there is an array of well characterised pharmaceutically available drugs that can be used to treat many diseases. The ability to identify and repurpose drugs is therefore a viable and cost effective means of restricting PyV-associated diseases.

The entry of Bunyaviruses and Filoviruses have been shown to require K^+^ and Ca^2+^ channels, respectively (38, 42). Therefore TEA and verapamil were applied during attachment and entry of MCPyV and SV40. The results indicated that MCPyV required both K^+^ and Ca^2+^ channel activity, whilst SV40 trafficking was solely sensitive to Ca^2+^ channel blockage (**Fig. 3**). The use of a wider panel of K^+^ channel inhibitors suggested that MCPyV required the activity of K_V_ channels (**Fig. 4A**). Insensitivity to K^+^ channel inhibitors during SV40 infection (**Fig. 4B**) highlighted mechanistic differences between MCPyV and SV40. Importantly, neither blocker inhibited SV40 so their effects cannot be mediated through modulation of endosomal pH. However, further screening with other human PyVs such as JCPyV and BKPyV may identify conserved requirements that could be targeted in the treatment of PyV-associated disease in humans.

Due to conserved sensitivity of MCPyV and SV40 upon challenge with verapamil, the role of Ca^2+^ channels was further explored. Treatment with flunarizine and nitrendipine, T- and L-type Ca^2+^ channel inhibitors, respectively, produced surprising results (**Fig. 5A-D**). Whilst MCPyV was sensitive to T-type Ca^2+^ channel inhibition there was no observed inhibition of SV40 with either drug, again highlighting differences between MCPyV and SV40 during entry. The lack of phenotypic change for SV40 was however comparable to data relating to Ebola virus (EBOV), where verapamil was shown to prevent docking of virion-containing endosomes with the ER through inhibition of NAADP-sensitive Ca^2+^ channels (41). EBOV inhibition was further characterised to be via NAADP-sensitive TPC1/2 activity, therefore we similarly investigated whether TPC inhibition could restrict MCPyV and SV40. Treatment with the L-type Ca^2+^ channel gabapentin had no effect upon entry of MCPyV or SV40 (**Fig. 6A&C**). However, treatment with the NAADP-sensitive Ca^2+^ channel inhibitor tetrandrine showed a concentration-dependent effect upon both viruses, with ablation of entry for MCPyV and SV40 at 5 µM and 10 µM, respectively (**Fig. 6B&D**). As tetrandrine-mediated inhibition of EBOV entry was due to the prevention of ER docking, a viral uncoating assay for SV40 was performed as further validation (**Fig. 7**). Consistent with previous results, following treatment with 10 µM tetrandrine VP2/3 was undetectable, confirming that virions either did not enter the ER or were unable to disassemble to reveal VP2/3 (summarised in **Fig. 8**). Identification that PyVs share a conserved requirement with EBOV suggests that NAADP-sensitive TPC inhibition represents a therapeutic target for viruses and pathogens that traffic through the ER during entry stages. Although tetrandrine is not widely available and there are currently limited studies into the efficacy of treatment *in vivo*, the identification of endosomal-ER fusion as a requirement for a variety of pathogens provides a common target that could potentially be exploited (60, 61).

**Figure 8:**
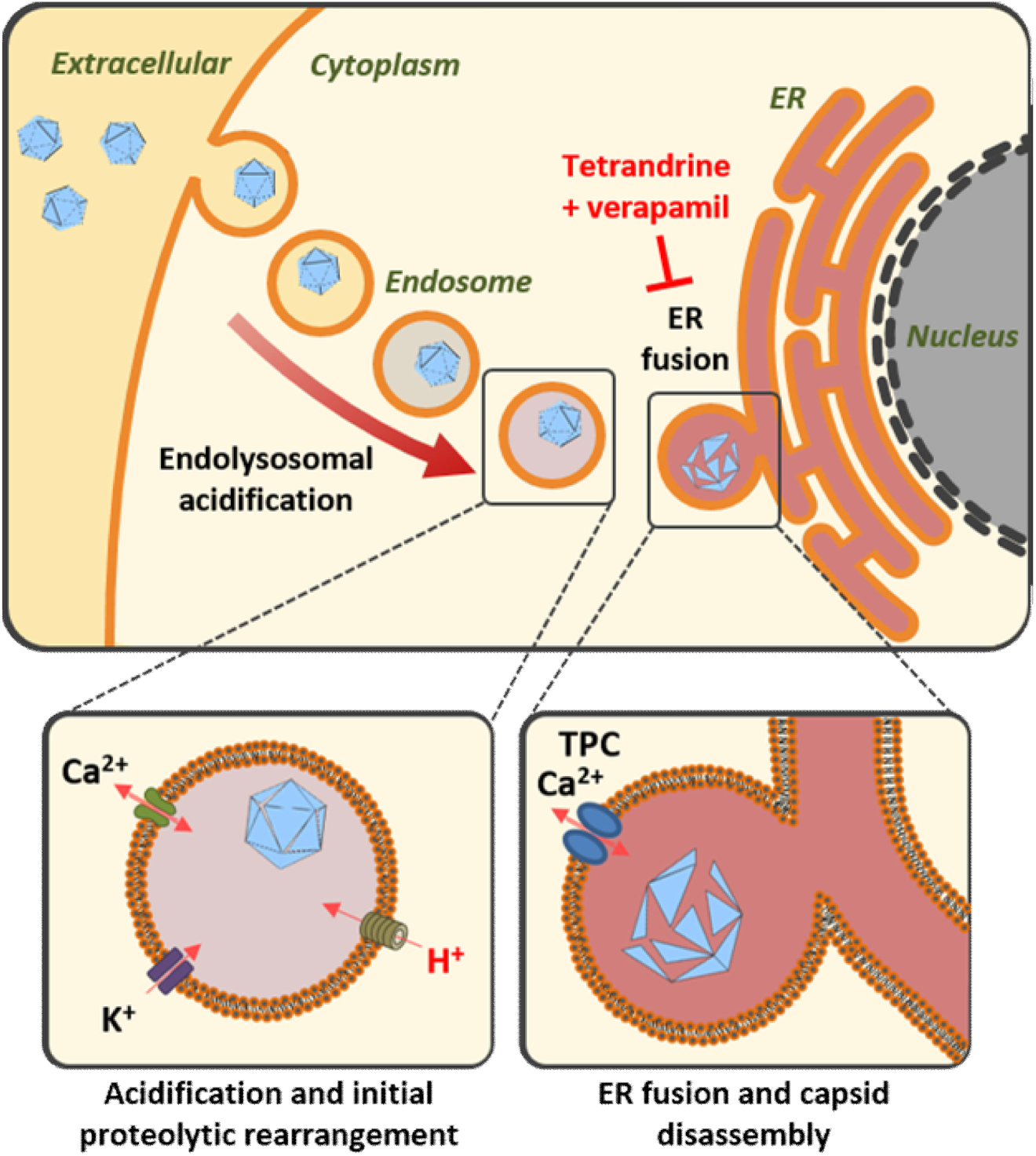
Schematic representation of proposed ion channel requirements during MCPyV and SV40 entry. Following internalisation, MCPyV and SV40 require an acidic environment. In addition to lowered pH, MCPyV also requires the activity of K^+^ and T-type Ca^2+^ channels whilst trafficking to the ER (**A**). Fusion to the ER of both MCPyV and SV40 is dependent upon NAADP-sensitive Ca^2+^ channel activity where the capsid disassembles, exposing the minor capsid proteins VP2/3 prior to translocation into the cytoplasm, which is inhibited by verapamil and tetrandrine.

In conclusion, we provide the first evidence that repurposing of clinically available drugs that modulate ion channel activity are a viable method of restricting PyV infection. This study identifies that MCPyV is more sensitive to channel inhibition than SV40, with K_V_ and T-type Ca^2+^ channel inhibition restricting entry which may be applicable to other humans PyVs. Additionally we have demonstrated that the NAADP-sensitive TPC inhibitor tetrandrine is a potent inhibitor of both MCPyV and SV40. Ca^2+^ channel modulation is therefore a potential mechanism through which human PyV diseases associated with persistent infection could be modulated. Coupled with previous studies, this requirement reveals a conserved target to restrict a wider range of pathogens that transit through the ER.

## MATERIALS AND METHODS

### Antibodies and chemicals

The (pAb)108 hybridoma used to detect SV40 T-antigens was a kind gift from Daniel DiMaio (Yale Cancer Centre, Connecticut, USA). SV40 VP2/3 antibodies were purchased from Abcam. Calnexin antibodies were purchased from Thermo Fisher Scientific.

EGA, gabapentin, KCl, quinine, TEA, tetrandrine and verapamil were purchased from Sigma-Aldrich. 4AP and flunarizine were purchased from Alfa Aesar. Nitrepndipine and NH_4_Cl were purchased from Santa Cruz Biotech.

### Cell lines and maintenance

HEK293TT cells were a kind gift from Christopher Buck (NIH, National Cancer Institute, Bethesda, MD, USA). Vero cells were a kind gift from Andrew Macdonald (University of Leeds, Leeds, UK). Cells were maintained using Dulbecco’s modified Eagle’s medium (DMEM) containing 10% (v/v) foetal bovine serum (FBS) and 50 U/mL penicillin and streptomycin (complete DMEM). HEK293TT medium was supplemented with 250 µg hygromycin B (Thermo Fisher Scientific) to maintain T-antigen expression, with removal prior to experimentation.

### SV40 production and titration

Stock of SV40 supernatants were provided by Andrew Macdonald (University of Leeds, Leeds, UK). Virus stocks were produced through infection of naïve Vero cells, with virus progeny containing medium removed 7 days post-infection. Supernatants were centrifuged at 16,000 *g* to pellet cellular debris and aspirated medium flash frozen and stored at −80 °C.

To determine virus stocks 5×10^3^ Vero cells were seeded into wells of a 96-well plate 18 hours prior to infection. Virus stocks were diluted 5-fold prior to a 2-fold dilution series using complete DMEM. Virus dilutions were incubated on cells in triplicate for 2 hours before aspiration and addition of fresh complete DMEM. Cells were fixed 24 hpi with 4% (w/v) paraformaldehyde (Sigma-Aldrich) and permeabilised in 0.1% (v/v) Triton X-100 (Fisher Scientific). SV40 T-antigens were detected using (pAb)108 and species-specific AlexaFluor 488 (Life Technologies, Thermo Fisher Scientific) antibodies. Wells were imaged using an Incucyte ZOOM instrument, with 4 non-overlapping images taken in each well. The number of T-antigen positive cells were counted per well. Reciprocals were calculated to identify dilutions with a linear relationship between dilution and T-antigen positive cells. Values were used to calculate IU/mL.

### SV40 infection assay

Vero cells (5×10^3^/well) were seeded into 96-well plate 18 hours prior to experimentation. If applicable, cells were pre-treated with chemical inhibitors for 1 hour. SV40 virions were diluted in complete DMEM (containing chemical inhibitors if applicable) to achieve an MOI of 1 relative to initial seeding and incubated on cells for 2 hours with gentle rocking every 30 minutes before aspiration and addition of fresh complete DMEM. Cells were fixed 24 hpi, immunostained, imaged and analysed using an Incucyte ZOOM System as previously described. Inhibitor effects were calculated through comparison to untreated controls. Percentage confluence was calculated by Incucyte ZOOM analysis to ensure cell proliferation in drug-treated wells, with <80% confluence omitted.

### MCPyV PsV production

Production of MCPyV PsVs has been previously described (14, 21, 62, 63). Briefly, 293TT cells were transfected with pwM2m, ph2m and pEGFP-C1 and harvested by trypsinisation 48 hours post transfection. Cells were pelleted before lysis using 0.5% Triton X-100 and incubated for 24 hours at 37 °C following addition of 1/1000^th^ volume RNase Cocktail enzyme mix (Ambion) and 25 mM ammonium sulphate (Sigma-Aldrich), pH 9.0. PsVs were extracted by centrifugation and loaded onto a 27-33-39% discontinuous opti-Prep (Sigma-Aldrich) gradients prior to ultracentrifugation. Fractions were collected and samples were analysed by Western blotting and silver staining to identify PsV-containing fractions, which were pooled and flash frozen prior to storage at −80°C. BSA standards were separated by SDS-PAGE alongside gradient fractions prior to silver staining to determine relative mass of PsVs in each fraction.

### MCPyV reporter assays

5×10^4^ 293TT cells were seeded into wells of a poly-L-lysine treated 24-well plate 18 hours before addition of PsVs. If required, pre-treatment with chemical inhibitors was performed for 1 hour. 10 ng VP1 equivalent of PsV stock was mixed with complete DMEM (containing chemical inhibitor if required) and added to wells for 2 hours with gentle shaking every 30 minutes before aspiration and addition of fresh complete DMEM. 72 hpt detection of GFP positive cells was performed using an Incucyte ZOOM System as previously described. Chemical inhibitor effects were calculated through comparison to an untreated control. Percentage confluence was calculated by Incucyte ZOOM analysis to ensure continued proliferation in comparison to untreated cells, with chemical inhibitors with <80% comparable confluence omitted.

### Minor capsid protein exposure assay

5×10^4^ Vero cells were seeded onto coverslips in 24-well plates 18 hours before experimentation. Plates were chilled at 4 °C for 30 minutes before addition of SV40 virions at an MOI of 3 in pre-chilled complete DMEM (containing inhibitors if applicable). Cells were kept at 4 °C for 1 hour with gentle agitation every 15 minutes to permit virus binding. Medium was then aspirated and replaced with pre-warmed complete DMEM (containing inhibitors if applicable) to synchronise infection. Cells were fixed 10 hours post infection and immunofluorescence performed as previously described (64). VP2/3 specific antibodies were used to detect exposed minor capsid proteins, with calnexin antibodies used to visualise proximity to the ER. Microscopy was performed using a ZEISS LSM 880 confocal microscope.

## ACKNOWLEDGEMENTS

The authors would like to thank Daniel DiMaio, Christopher Buck and Andrew Macdonald for kindly providing reagents used in this study. We are grateful to members of the Whitehouse laboratory for helpful discussions. The work was funded in parts by a MRC studentship (95505126) and Royal Society University Research Fellowship to JM (UF100419).

